# Identification of Diverse Cellulose Binding Domains using *in silico* Prioritisation and High-Throughput Screening

**DOI:** 10.1101/2025.10.06.677802

**Authors:** Christopher R. Field, Ashley P. Mattey, Sebastian C. Cosgrove, Sam Hay

## Abstract

*In silico* pre-screening methods are becoming an increasingly important step in making large scale enzyme screening experiments manageable, particularly in the sampling large protein sequence datasets for maximum diversity. Here, we develop a bioinformatics workflow that utilises automated gene sequence annotation, substrate prediction and structure prediction tools to effectively isolate representative protein sequences, thereby maximising the information gathered on the wider dataset from relatively few *in vitro* experiments. Included in this workflow is a new tool, ColabAlign, which performs pairwise structural alignments to construct a structure-informed dendrogram, from which representatives are selected based on clustering. The workflow was applied to the identification of cellulose-binding carbohydrate-binding modules (CBMs) suitable for enzyme immobilisation tag development. 47 sequentially and structurally diverse mCherry-CBM fusions were tested for binding against cellulose, chitin and spent coffee grounds (SCG) using a pulldown assay from cell lysate. We successfully identified 5 CBMs with significant binding towards commercial and waste cellulose support materials, suitable for further tag design work. This work also provides the first experimental evidence of chitin binding by one or more members of CBM6 and CBM46.

## Introduction

As datasets of genes encoding putative proteins continue to grow, even the most high-throughput screening and characterisation methods require *in silico* methods for selection and prioritisation prior to experimental testing. Given that sequence data alone does not provide complete information about a given protein and those related to it, a number of *in silico* screening methods have emerged to predict aspects such as enzyme-substrate pairs (1,2), soluble expression (3,4) and inhibitor binding through molecular docking (5,6). These layers provide a level of pre-selection meaning, ideally, only active proteins are expressed, improving the quality received from a relatively small number of experiments.

One area with a large dataset of uncharacterised proteins is carbohydrate-binding modules (CBMs). CBMs, previously cellulose-binding domains (CBDs), are a class of non-catalytic protein domains found appended to carbohydrate-active enzymes (CAZymes). They significantly improve the efficiency of CAZyme-mediated hydrolysis of oligo- and polysaccharides through the combination of targeting and proximity effects, *i*.*e*., directing the associated CAZyme to the substrate, and increasing local CAZyme concentration by limiting CAZyme dissociation, respectively (7). This is also thought to allow processive CAZyme action, enabling consecutive rounds of hydrolysis on long carbohydrate chains (7-9). CBM-polysaccharide binding is thought to involve a combination of hydrogen bonding with proximal polar residues, and relatively weak CH-π interactions with aromatic residues (10,11), the most energetically favourable of which being tryptophan (12).

CBMs are a ubiquitous type of protein domain. Since their first identification in fungal species *Trichoderma reesei* (13), they have been putatively or experimentally identified across all kingdoms, as well as in viruses (14). CBMs are primarily categorised by sequence similarity, with 107 distinct sequence families identified to date. These families cluster CBM-containing sequences from the over 5.6 million entries currently listed in the Carbohydrate-Active enZYmes (CAZy) database (15). Sequence families exhibit unique substrate binding capabilities, ranging from large, insoluble polysaccharides (16,17), to shorter mono- and oligo-saccharides (18-21), and even synthetic polymers including polyethylene terephthalate (PET) and polystyrene (PS) (22,23). Such variations are, in part, a product of the three different methods by which CBMs bind their substrates: Type A/*planar* CBMs feature a flat binding surface with exposed aromatic residues that binds crystalline polysaccharide regions; Type B/*exo* CBMs typically feature a binding cleft with aromatic residues positioned above and below isolated oligosaccharide chains; Type C/*endo* CBMs feature smaller binding pockets that encapsulate a single, terminal monosaccharide unit (24). CBMs can also be categorised into 1 of 7 secondary structural fold families (**Figure 1**), the most common of these motifs being a β-sandwich (fold family 1) which consists of two β-sheets typically composed of between 3 and 6 antiparallel β-strands (24-26).

**Figure 1.**
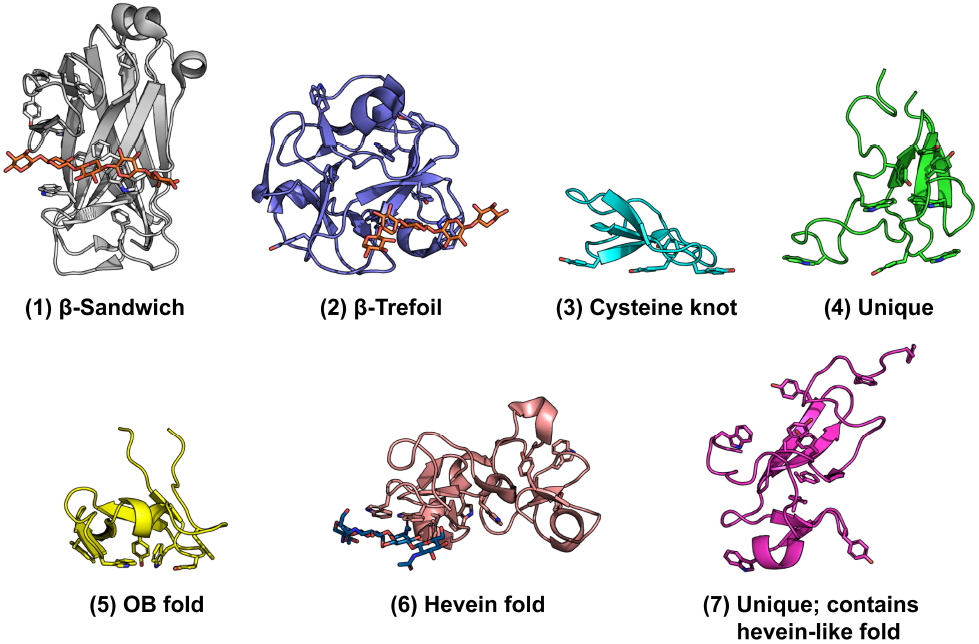
Example X-ray/NMR structures of the 7 CBM fold families, as defined by Boraston et al. (25). Folds are rendered from PDB IDs: 1GNY, 1MC9, 1CBH, 1AIW, 1E8R, 1EN2 and 1DQC, respectively (27-33). Aromatic residue side chains and, where present, bound oligosaccharides are rendered as sticks. Ligands are coloured following SNFG notation (34): blue, glucose-derived oligosaccharides; orange, xylose-derived oligosaccharides.

In recent years, CBMs have emerged as a promising, low-cost tool for many biotechnological applications, including protein purification (35,36), bioethanol production (37), antibody immobilisation (38) and enhanced enzymatic plastic degradation (22,23). One area receiving particular focus is their use as a relatively green method of immobilising enzymes on polysaccharide support materials (39-43). Enzyme immobilisation provides several benefits over free enzyme processes, namely improved enzyme stability over long reaction runtimes and simplified catalyst-product separation (44,45). Further, it enables enzymatic reactions to be integrated into flow chemistry systems, which provide additional benefits over batch reactions including improved atom economy through cofactor recycling (46,47), mitigation of substrate inhibition (48), inline product analysis (49), improved safety by reducing reactor sizes (50), and the ability to run multi-enzyme cascade reactions with each enzyme under discrete conditions (45,51). To date, the scope of CBM-enzyme immobilisation studies has been limited to a relatively small number of CBMs already well characterised in the literature. Despite recent high-throughput screening efforts (23,52), the majority of the CBM sequence space remains unmapped, particularly within the context of immobilisation tag design.

To address this gap in knowledge, and to identify promising CBM candidates for further development as protein/enzyme immobilisation ‘tags’, we developed a bioinformatics workflow that facilitates broad screening of large protein datasets, maximising the information gathered from a small number of *in vitro* experiments. Included in this workflow is a new tool, ColabAlign (53), which constructs dendrograms based on the structural similarity of experimentally resolved and predicted protein structures, and selects representative proteins using clustering and structure-informed sequence alignment. We used this workflow to select 47 maximally diverse CBMs, covering 17 sequence families and expressed them as fusion proteins with mCherry. The mCherry-CBM fusions where then screened *in vitro* using a pulldown assay from cell lysate. CBMs were tested for binding against a commercially available chitin resin and 2 microcrystalline cellulose powders, with average particle sizes of 50 μm and 90 μm, to investigate the impact of additional functional groups and particle size/morphology on CBM-polysaccharide binding. This is pertinent to their translation to continuous flow biocatalysis systems, where the physical properties of support material can have a substantial impact on pressure drop across a packed-bed reactor (54). Finally, binding to spent coffee grounds (SCG) was also investigated as SGC represents an abundant (∼6 million tonnes is produced annually (55)) and sustainable support material for CBM-mediated enzyme immobilisation, due to its high cellulose and hemicellulose content, which is ∼50% of dry weight (56).

## Materials and Methods

### Reagents

Polymyxin B sulphate (catalogue # BIP0145) was purchased from *Apollo Scientific*; Kanamycin sulfate (catalogue # M02038) was purchased from *Fluorochem*; Auto-induction lysogeny broth base with trace elements (AI-LB) (catalogue # AIMLB0205) was purchased from *Formedium*; Avicel^®^ PH-101 (50 µm average particle size) (catalogue # 11365), Deoxyribonuclease I from bovine pancreas (DNase I) (catalogue # DN25) and Lysozyme from chicken egg white (catalogue # L3790) were purchased from *Merck*; Chitin resin (catalogue # S6651S) was purchased from *New England Biolabs*; 90 µm average microcrystalline cellulose (catalogue # 382312500), Thermo MCC, was purchased from *Thermo Fisher Scientific*; SCG was generated in-house from single origin Guatemala coffee used to prepare espresso.

### Biological Resources

mCherry-CBM constructs were synthesised and inserted into the pET-28a(+) expression vector between the *NcoI* and *XhoI* restriction sites by *Twist Bioscience* (San Francisco). A flexible Gly-/Ser-rich linker designed by Waldo et al. (57), GSAGSAAGSGEF, was inserted between C-terminus of the mCherry and N-terminus of each CBM. The construct was positioned in frame of the hexa-His-tag present directly 3’ of the *XhoI* site. An example plasmid map is given in **Figure S1**. Chemically competent *Escherichia coli* strains *BL21(DE3)* (catalogue # C2527H) and *SHuffle T7* (catalogue # C3026J) were purchased from *New England Biolabs*.

### Recombinant protein expression

mCherry-CBM constructs were transformed into strains *BL21(DE3)* or *SHuffle T7* cells depending on the predicted presence of internal disulfide bonds. CBMs were classified as possibly containing one or more internal disulfide bonds if they contained at least 2 cysteine residues with sulfur atoms within 5 Å. This distance was notably longer than the measured sulfur-sulfur distance in a disulfide bond (2.03 Å) (58) to account for unfavourable cysteine site chain positioning in the AlphaFold models. Cultures expressing each mCherry-CBM fusion were grown in triplicate from glycerol stocks in 24-well deep well plates. Cultures were grown in 2 ml AI-LB containing 25 µg/ml kanamycin at 37°C (*BL21(DE3)*) or 30°C (*Shuffle T7*), 850 RPM, 80% humidity for 4 hours. The temperature was then dropped to 25°C for a further 32 hours. Cell pellets were harvested by centrifugation at 4°C, 4,000 RPM for 20 mins and stored at -20°C until needed.

### mCherry-CBM pulldown assay

20 g of SCG was washed with 100 ml of diH_2_O prior to use, to minimise the impact of other compounds quenching mCherry excitation. 20 g of all 4 supports (Avicel PH-101, Thermo MCC, chitin resin, SCG) were then washed with 30 ml 1x phosphate-buffered saline (PBS, pH 7.4). Excess buffer was removed by centrifugation at 4,000 RPM for 2 mins. Cell pellets were resuspended in 2 ml lysis buffer (1x PBS pH 7.4; 1.0 mg/ml lysozyme; 5 µg/ml DNase I; 0.5 mg/ml polymyxin B) and incubated at 30°C, 850 RPM, 80% humidity for 2 hours. Cell debris was removed by centrifugation at 4°C, 4,000 RPM for 20 mins. 400 µl of cleared lysate was combined with 100 mg of each support and incubated at 30°C, 850 RPM, 80% humidity for 1 hour. Support materials were separated from the post-binding lysate by centrifugation at 4°C, 4,000 RPM for 10 mins. 100 µl of cleared cell lysate at post-binding lysate were transferred to a black 96 well plate, and fluorescence was measured using a BMG Labtech FLUOstar Omega, with an excitation filter of 584 nm and emission filter of 610-620 nm. Binding Intensity (BI) values were determined using Eq. 1:

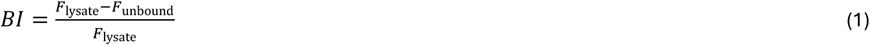

where *F* is fluorescent intensity. mCherry-CBM fusion concentrations were determined by interpolation of an mCherry standard curve, which was quantified based on His-mCherry concentration determined by ε(280 nm) = 34,380 M^-1^ cm^-1^, as calculated using *Expasy* ProtParam (59).

### Initial CBM sequence identification

The initial list of gene sequences was obtained from CAZy (15) on February 26^th^ 2024, which contained a total of 4,448,707 GenBank code entries. This included 497,515 entries annotated as containing at least 1 CBM-encoding region. Duplicate codes were filtered, leaving 376,307 unique GenBank codes: 333,309 from the *National Center for Biotechnology Information* (NCBI) (60) and 42,998 from *Joint Genome Institute* (JGI) (61). JGI sequences were excluded from the dataset due to difficulties in retrieving sequences in bulk. Sequences for the NCBI CBM entries were downloaded and duplicate CBM-containing sequences were filtered using Biopython 1.83 (62), leaving 197,726 unique genes containing at least 1 CBM-encoding region.

## Results/Discussion

An overview of the bioinformatics workflow shown in **Figure 2**. Initially, a library of 197,726 unique CBM-containing gene sequences was assembled. This contained a mixture of sequences encoding one or more identical or different CBM, as well as CAZyme domains. To isolate unique CBM-encoding sequences, we performed automated CAZy gene annotation using dbCAN3 (63). First, CBM-encoding regions were identified using the HMMER, dbCAN-sub and DIAMOND annotation tools. CBM regions identified by none or 1 of the tools were discarded to ensure accurate annotation. Gene start and end positions for the CBM-encoding regions were merged from the HMMER and dbCAN-sub outputs since the DIAMOND output only reported the length of the input gene. In cases where start and end values differed between the outputs, but where the ranges overlapped, the lower of the two start values and higher of the two end values were chosen to maximise gene coverage. These regions were written to a single FASTA-formatted file, and duplicate sequences were removed, leaving 158,459 unique CBM-encoding sequences originating from 123,312 unique GenBank entries.

**Figure 2.**
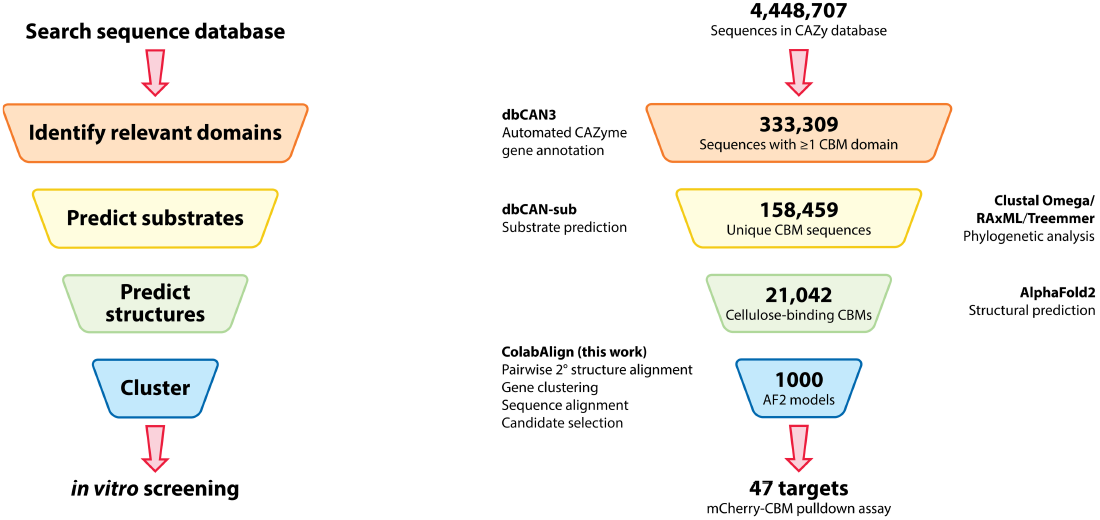
Visual summary of bioinformatics workflow. *Left:* Generalised workflow. *Right:* Workflow specific to this work, highlighting the number of CBM sequences at each stage, the tools used and their functions.

### *In silico* candidate selection

To avoid mistakenly including CBMs from CAZyme-CBM fusions where only the CAZyme was predicted to bind cellulose, the isolated CBM sequences were processed with dbCAN-sub a second time. The output was parsed for entries predicted to bind cellulose based on their assigned subfamily, reducing the dataset size to 21,042 sequences. Near-identical sequences were filtered from the dataset by clustering sequences of the same length, identifying groups within each cluster where sequences had 3 or few mismatched residues, and selecting a representative sequence for each group at random to improve the performance of phylogenetic tree generation. This reduced the dataset to 17,590 sequences. To group together sequentially similar CBMs, a global multiple sequence alignment was calculated using Clustal Omega (64), and a phylogenetic tree was generated using RAxML-NG (65) with the BLOSUM62 amino acid substitution matrix. The number of taxa on the phylogenetic tree was iteratively trimmed to 1000 sequences using Treemmer (66), maintaining as much of the sequence diversity of the original tree as possible.

To maintain the structural diversity of the reduced dataset, candidates from the remaining 1000 sequences were selected based on predicted structures generated with AlphaFold version 2.1.1 (67). Structures with a predicted local distance difference test (pLDDT) score of less than 90 were discarded to ensure accurate structural alignments were possible. After the first round of structure prediction, 66 models did not reach the pLDDT > 90 threshold, either because these sequences confer unknown protein folds, or more likely, the CBM-encoding regions determined by dbCAN3 did not match the true CBM-encoding region. Sequences associated with these models were extended at both termini using up to 5 residues from their original sequences. Following the second round of structure prediction, 61 of the original 1000 sequences did not meet the threshold and were discarded, leaving 939 CBMs in the dataset.

A maximally diverse subset of CBMs were selected from the AlphaFold models using ColabAlign v 0.1.0 (53). ColabAlign is a script that combines the functions of several existing bioinformatics tools to compare the structural similarity of a set of proteins using all-against-all pairwise alignment and select representative structures from the input set using clustering and structure-informed sequence alignment (**Figure 3**). The script utilised multiprocessing to run several instances of single-threaded programs in parallel, thereby considerably speeding up the processing time of datasets containing hundreds of structures. The tools used as part of ColabAlign include US-align (68) for structural alignment; Biopython (62) for dendrogram construction using the UPGMA algorithm (69); TreeCluster (70) for dendrogram clustering; MUSTANG (71) for calculating sequence alignments based on structural alignments; and MView (72) for calculating cluster consensus sequences and selecting candidates based on sequence similarity to the 60% identity consensus. Additional information on ColabAlign is presented in the Supporting Information.

**Figure 3.**
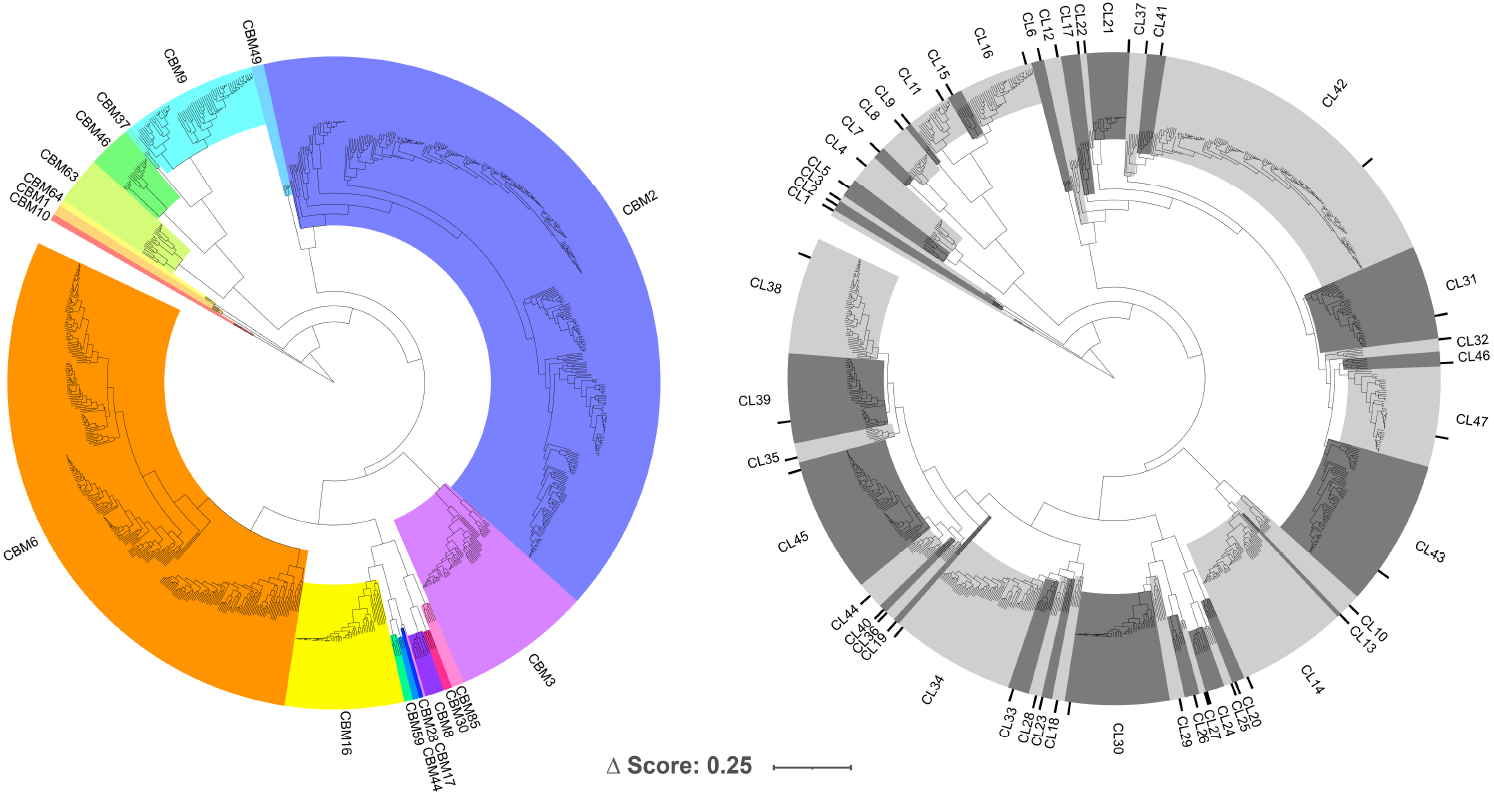
Structural alignment dendrogram from the ColabAlign output. Tree annotations were made using Interactive Tree of Life (iTOL) (117). *Left:* CBM sequence families separated by colour. *Right:* Clusters (CL) of structurally similar models. Representative CBMs from each cluster selected for *in vitro* characterisation are highlighted with black bars.

A total of 47 CBMs were selected from the initial dataset as representative genes (**Table 1**), predicted to be capable of binding at least one form of cellulose. A complete list of GenBank accession codes, determined CBM-encoding region, organism of origin, expression strain and predicted substrate binding capabilities is given in **Table S1**. The CBM sequence found in GenBank entry AFI25187 was originally annotated as both a member of family 17 and 28, due to the high structural similarities between the families (**Figure 3**; *left*), and sequence conservation suggesting a common ancestral CBM sequence (73). Structural alignments with published crystal structures from both families revealed a significantly higher global similarity between AFI25187 and the CBM28 crystal structures (mean TM-score 0.94 ± 0.02; *n* = 5) as opposed to the CBM17 structures (mean TM-score 0.70 ± 0.01; *n* = 2) (**Figure S2**). This CBM was therefore categorised as a family 28 CBM, and herein denoted *Ha*CBM28-CL27.

**Table 1.** Summary of selected CBM candidates.^a^.

| **Family** | **n** | **Fold** | **BM** | **Known family binding affinities** |  |  |
| --- | --- | --- | --- | --- | --- | --- |
|  |  |  |  | **Cellulose** | **Chitin** | **Hemicelluloses** |
| 1 | 1 | 3 | A | crystalline (16,74,75) | (74) | - |
| 2 | 12 | 1 | A | crystalline (76-79)<br>amorphous (77,78) | (80) | xylan (81) |
| 3 | 3 | 1 | A | crystalline (10,82-88)<br>amorphous (82,83,85,86,88) | (88,89) | xylan (89)<br>xyloglucan (90) |
| 6 | 13 | 1 | B | crystalline (91)<br>amorphous (91) | - | arabinoxylan (92)<br>mannan (52)<br>MLG (52,93)<br>xylan (91,92) |
| 8 | 1 | 1 | A/B | crystalline (94) | - | xyloglucan (94)<br>glucomannan (94) |
| 9 | 3 | 1 | C | crystalline (95)<br>amorphous (95)<br>short oligos (95) | - | xylan (95,96)<br>xyloglucan (95)<br>arabinoxylan (97)<br>mannan (96) |
| 10 | 1 | 5 | A | crystalline (98,99) | - | - |
| 16 | 2 | 1 | B | amorphous (100) | - | glucomannan (100) |
| 17/28 | 1 | 1 | B | amorphous (18,101-104) | - | MLG (18,101) |
| 30 | 1 | 1 | A/B | amorphous (105,106) | - | MLG (106)<br>xyloglucan (106) |
| 37 | 1 | 1* | A/B | crystalline (107)<br>amorphous (107) | (107) | MLG (107)<br>xylan (107) |
| 44 | 1 | 1 | B | crystalline (106)<br>amorphous (106) | - | MLG (106)<br>xyloglucan (106)<br>glucomannan (106) |
| 46 | 2 | 1 | A | crystalline (108,109) | - | MLG (109)<br>xyloglucan (109) |
| 49 | 1 | 1 | A | crystalline (110) | - | - |
| 63 | 2 | 1 | A/B | crystalline (111,112) | - | arabinoxylan (111) |
| 64 | 1 | 1 | A | crystalline (113) | (113) | - |
| 85 | 1 | 1* | A/B | crystalline (114)<br>amorphous (114) | - | glucomannan (114)<br>glucuronoxylan (114)<br>MLG (114) |

<sup>a</sup> n, number of candidates from a given family selected for *in vitro* screening; Fold, fold family numbers as defined by Boraston et al. (25): Fold family 1 is β-sandwich; 3 is cysteine knot; 5 is OB-fold; \*, predicted fold family based on AlphaFold structures where no crystal structures currently exist for the family; BM, binding mode; Known binding affinities were determined with the help of CAZypedia (26), and the CAZy (15), dbCAN-sub (115) and InterPro (116) databases, in addition to the references in the Table.

### mCherry-CBM Pulldown Assay

The 47 CBMs summarised in **Table 1** were expressed in *E. coli* as fusion proteins with an N-terminal mCherry domain to monitor expression and binding to support materials. Protein concentration was estimated using the fluorescence of His-tagged mCherry (i.e., without a CBM domain) as a reference. Change (loss) in fluorescence after pulldown by the substrate, expressed as mean BI (Eq. 1), was then used as a semi-quantitative metric of binding/capture (**Figure 4**). Blank-corrected fluorescence data are given in **Dataset S1**. Mean BI values for His-mCherry were 0.11 ± 0.04 for Avicel, 0.11 ± 0.05 for Thermo MCC, 0.23 ± 0.07 for chitin, and 0.14 ± 0.13 for SCG. All 47 mCherry-CBM fusions had BI values above the mCherry control for Avicel and Thermo MCC,40 for chitin and 42 for SCG. From these data, *Cc*CBM2-CL42, *Ma*CBM2-CL43, *Sc*CBM2-CL46 and *Cs*CBM64-CL3 had BI values above 0.7 for Avicel and Thermo MCC binding, making them the most promising candidates for future CBM immobilisation tag design for cellulosic support materials. These data also provide, to the best of our knowledge, the first experimental evidence of chitin binding by one or more members of CBM6 (13/15 above BI = 0.23) and CBM46 (2/2 above BI = 0.23). In addition, *Cc*CBM2-CL42, *Ma*CBM2-CL43, *Sc*CBM2-CL46 and *Sa*CBM6-CL44 had BI values above 0.5 towards SCG, suggesting they may be good targets for engineering SCG binding.

**Figure 4.**
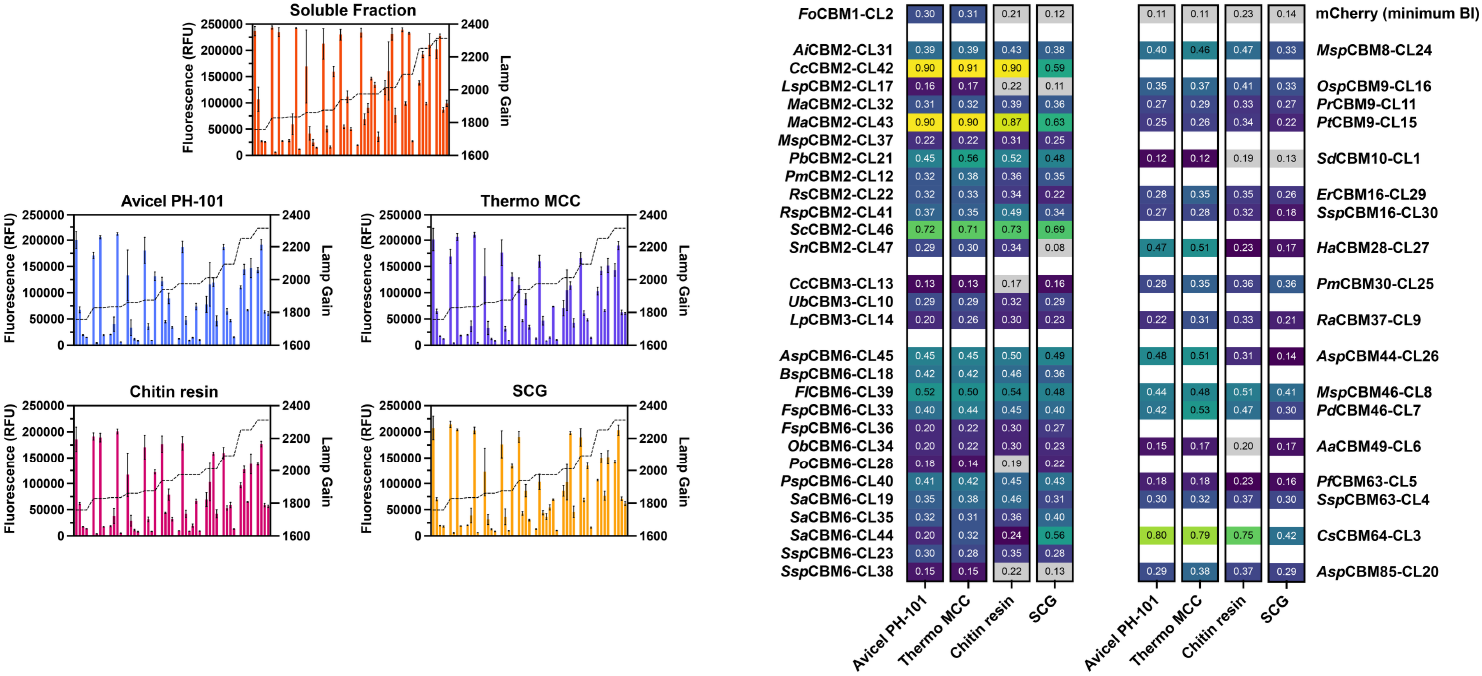
*Left:* Blank-corrected fluorescence values before (orange) and after incubating with Avicel PH-101 (blue), Thermo MCC (purple), chitin resin (magenta) and SCG (yellow). Dotted lines indicate plate reader lamp gain used to measure fluorescence per sample. *Right:* Mean BI values for mCherry-CBM binding grouped by sequence family. Values shaded in grey indicate values below the BI for the mCherry control.

To investigate specificity of the CBMs, a correlation analysis was performed. A strong linear correlation between BI values for Avicel and Thermo binding was observed (R^2^ = 0.91; **Figure 5**), indicating the differences in morphology between the two samples (*i*.*e*., particle size) has a minimal impact on binding. It is highly likely that the mCherry-CBM concentrations are not high enough to fully saturate the binding capacities of either Avicel or Thermo, at which point divergence in BI values resulting from morphological differences would likely become more apparent. Correlations between the cellulose and chitin samples were also high, with R^2^ = 0.81 and 0.82 for Avicel PH-101 and Thermo MCC, respectively. Chitin and cellulose are both β-1,4 linked polysaccharides, so some correlation is expected. However, some cases where chitin binding was markedly higher (e.g., *Rsp*CBM2-CL41 and *Sa*CBM6-CL19) or lower (e.g., *Ha*CBM28-CL21 and *Asp*CBM44-CL26) than cellulose binding were observed. Differences may arise when substrate binding to these CBMs includes specific interactions between the protein and the hydroxyl (cellulose) or acetylamino (chitin) moieties at the C2 position of the substrate (**Figure S3**). Binding towards SCG was on average lower than for the other support materials, which is reflected in the markedly lower R^2^ values for this support. This is to be expected as the total cellulose content per gram for SCG will have been lower than the pure cellulose supports. Interestingly, *Sa*CBM6-CL44 displayed increased binding towards SCG over other support materials, likely indicating it can bind to one or more hemicellulose. Current experimental findings for family 6 CBMs suggests either arabinoxylan, mannan, mixed-linkage glucan (MLG) or xylan (**Table 1**), however further characterisation is required to confirm this.

**Figure 5.**
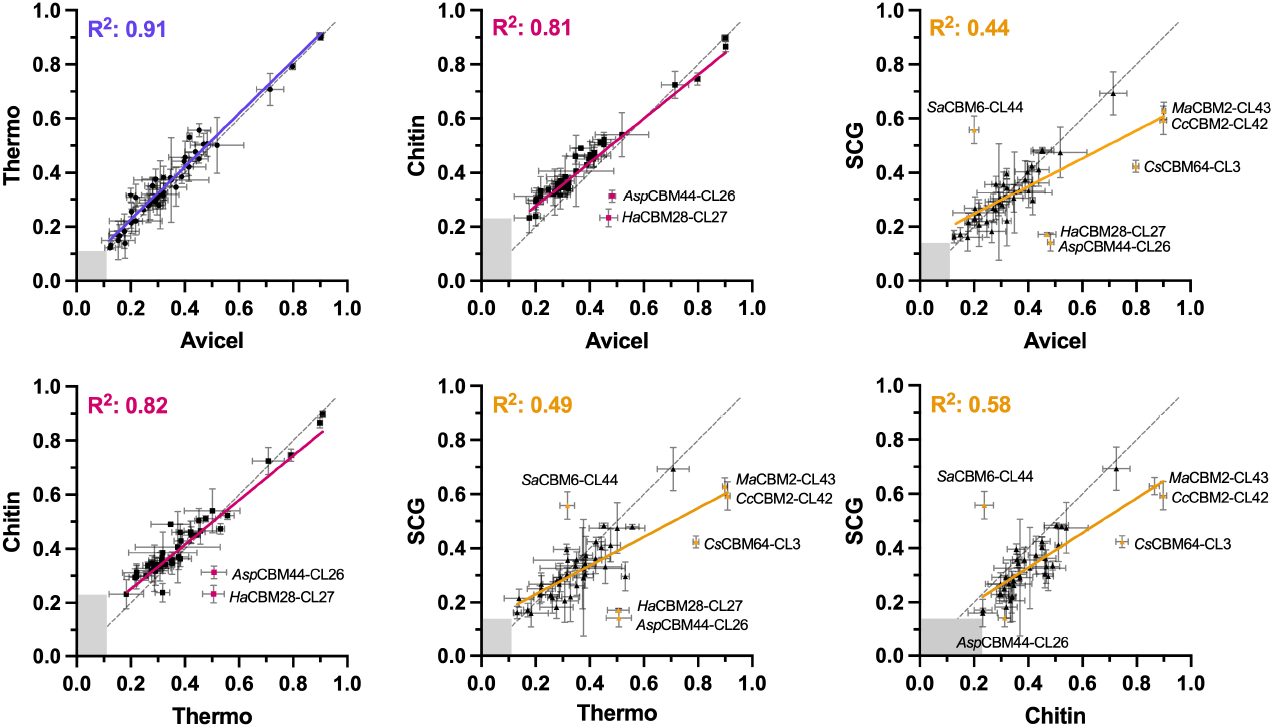
Correlation analysis comparing BI values for CBM binding to each combination of support materials. Names of significant outliers are listed next to their data. The black dashed line shows the diagonal, and shaded areas indicate the minimum required BI values to show binding above those of the His-mCherry control (Avicel: 0.11; Thermo MCC: 0.11; chitin: 0.23; SCG: 0.14).

### Comparisons with recent high-throughput CBM screening studies

Recent works by Salmeán et al. (52) and Rennison et al. (23) demonstrate alternative methods of performing high-throughput CBM characterisation screens. Salmeán et al. (52) utilised a double-blind comprehensive microarray polymer profiling (DB-CoMPP) method to screen 10 CBMs and 14 starch utilizing system protein D-like (SusD-like) proteins against a wide range of commercial polysaccharides and algal/plant cell wall extracts. Rennison et al. (23) utilised an automated holdup assay to successfully express and screen 797 type A CBMs from 4 families (CBM2, CBM3, CBM10 and CBM64) against a range of natural and synthetic polymers. Type A CBMs bind their substrates with a flat binding face to crystalline regions of their substrates, making them ideal for binding to large synthetic polymer samples (*e*.*g*., PET) (24). Further, examples of type A CBMs from these 4 families have been previously discovered with nanomolar affinities (*K*_d_) towards PET (118). While we screened substantially fewer CBMs (47 total), we covered a much wider range of families (17 families) and binding modes (type A, B and C). Consequently, our approach complements these studies and could be used as a pre-screen to identify new targets prior to a larger high-throughput screening. Also, we were able to assay the mCherry-CBM fusions directly from cleared cell lysate, following the same principle as previous one-step purification/immobilisation protocols (35,41,119). This allowed us to measure BI values for CBMs that had previously not been characterised due to low titres of purified protein. However, using cleared cell lysate with the holdup assay would likely not be possible without a more thorough lysate clearance step, as the filtered 384 well assay plates would become easily clogged with debris.

Directly comparing the results of the pulldown (this work) and holdup (23) assays reveals 7 CBMs present in both screens (**Table 2**), and some disagreement between our findings. Larger Avicel and chitin BI values were observed for *Pm*CBM2-CL12 in our pulldown assay than in the holdup assay. Conversely, *Lp*CBM3-CL14 and *Ub*CBM3-CL10 performed significantly worse in our pulldown assay. This may have resulted from slight differences gene sequence ranges used for these CBMs, arising from conflicting sequence annotations. Such differences can have a major impact on protein expression and folding, possibly explaining the observed differences in binding. Additionally, *Sc*CBM2-CL46 and *Sd*CBM10-CL1 both contain 2 cysteine residues with inter-sulfur distances of ∼2 Å in our AlphaFold models, indicating a high likelihood of cysteine bond formation. We therefore expressed these CBMs, in addition to those listed in **Table S1**, in *Shuffle T7* cells, which support cytosolic disulfide bond formation through a chromosomally-integrated disulfide bond isomerase (DsbC) (120). The lack of soluble expression seen in the holdup assay likely stems from misfolding due to *BL21(DE3)* lacking an equivalent endogenous enzyme. Whilst we were able to measure BI values for *Rs*CBM2-CL22 and *Sc*CBM2-CL46, the overall expression levels of these constructs were lower than many of the other mCherry-CBM fusions (**Table S3**), in agreement with the holdup results.

**Table 2.** Comparison of Avicel and chitin BI values for CBMs appearing in the pulldown (this work) and previous holdup (23) assays.^a^.

| **Name** | **GenBank** | **Support** | **Pulldown** | **Holdup (23)** | **$\Delta$ BI** |
| --- | --- | --- | --- | --- | --- |
| *Pm*CBM2-CL12 | ABM79533 | Avicel | 0.32 | NB | 0.19 |
|  |  | Chitin | 0.36 | 0.11 | 0.25 |
| *Rs*CBM2-CL22 | ABV54446 | Avicel | 0.32 | NE | - |
|  |  | Chitin | 0.34 | NE | - |
| *Sc*CBM2-CL46 | CAN95714 | Avicel | 0.72 | NE | - |
|  |  | Chitin | 0.73 | NE | - |
| *Cc*CBM3-CL13 | ADL50319 | Avicel | 0.13 | NB | 0.00 |
|  |  | Chitin | 0.17 | 0.11 | 0.06 |
| *Lp*CBM3-CL14 | ABX43721 | Avicel | 0.20 | 0.52 | 0.32 |
|  |  | Chitin | 0.30 | 0.30 | 0.00 |
| *Ub*CBM3-CL10 | AHL27899 | Avicel | 0.29 | 0.75 | 0.46 |
|  |  | Chitin | 0.32 | 0.54 | 0.22 |
| *Sd*CBM10-CL1 | ABD79918 | Avicel | 0.12 | NE | - |
|  |  | Chitin | 0.19 | NE | - |

<sup>a</sup> NB, no binding (*i.e.*, binding below untagged EGFP levels on that substrate); NE, no expression (*i.e.*, purified CBM-EGFP fusion titres did not exceed the 0.5 $\mu$ M required for the holdup assay). $\Delta$ BI values for NB samples were calculated assuming a maximum possible BI of 0.13 (mean Avicel-EGFP BI).

### *In silico* protein expression screening

Introducing a soluble expression prediction step into the pre-screening workflow (**Figure 2**; after predicting cellulose-binding CBMs) would provide an additional filter to remove proteins with low predicted soluble expression in *E. coli*, thereby maximising the number of successful assays. Sequences for the 47 selected CBMs were processed with 3 modern protein solubility prediction tools, SoluProt, (3), NetSolP (4) and DeepSoluE (121), to ascertain the relationship between predicted and observed soluble CBM fusion protein expression in *E. coli*. As the pulldown fluorescence values were measured on a per-plate basis to ensure accurate measurements were taken, these data could not be directly compared to estimate expression. We therefore used fluorescence values from a preceding expression trial (**Figure S4**), which were measured using identical gain values. From these data, we found no correlation between the scores of the three solubility prediction tools and the measured fluorescence values (max R^2^ = 0.05; **Figure S5**). To confirm whether this was caused by a small sample size, each method was repeated using purified EGFP-CBM concentration data provided by Rennison et al. (23). These data produced a similar absence of correlation between predicted and measured protein expression (max R^2^ = 0.02; **Figure S5**), leading us to conclude that these tools may not be suitable for predicting CBM or CBM fusion protein expression in *E. coli*.

### Challenges involved in creating a sequence-function relationship for CBMs

Experimental evidence of binding to one or more forms of cellulose exists for at least 15 families not picked up by dbCAN-sub (**Table S2**). This is symptomatic of the challenges in developing a sequence-function relationship for CBMs. Since CBMs are non-catalytic proteins, no clear method exists for classifying their function, unlike the Enzyme Commission (EC) number used to classify catalytic proteins. This limits tools such as dbCAN-sub to stochastic modelling techniques based on incomplete experimental data, which, in turn, restricts the scope of targeted CBM engineering studies. This exemplifies the need for both high-throughput screening data, such as the work by Salmeán et al. (52) and Rennison et al. (23), and a breadth of sequentially and structurally diverse CBMs to be characterised, to better cover the CBM sequence and structure spaces and make developing a robust sequence-function relationship possible. Such a relationship has the potential to improve our ability to predict the substrate binding preferences for uncharacterised CBMs from existing or newly established families, and to dramatically accelerate the speed at which CBM can be evolved or engineered towards unnatural substrates, notably synthetic polymers (e.g., plastics, epoxy resins), with the goal of significantly improving enzymatic breakdown of plastic through the addition of bespoke engineered binding tags.

## Conclusions

We have established a combined bioinformatics and high-throughput screening workflow that enables the characterisation of a maximally diverse set of proteins. We applied this workflow to select 47 CBMs from an initial dataset of over 4 million sequences, which were successfully expressed as fusion proteins with mCherry and assayed for binding to cellulose, chitin and SCG. We identified *Cc*CBM2-CL42, *Ma*CBM2-CL43, *Sc*CBM2-CL46, *Sa*CBM6-CL44 and *Cs*CBM64-CL3 as candidates for further CBM-based enzyme immobilisation tag design for commercial and waste cellulosic support materials. Our data also provides the first experimental evidence of CBM6 and CBM46 binding to chitin. The workflow can be adapted to different enzyme classes by swapping out CBM-specific tools for tools relevant to the target proteins. It can also be scaled up to assist in screening a larger, but still diverse set of CBMs using an automated high-throughput screening protocol.

## Data Availability

Source code for ColabAlign is available on GitHub under the MIT licence (https://github.com/crfield18/ColabAlign) and Zenodo (https://zenodo.org/records/14169501).

## Acknowledgements

The authors would like to acknowledge the assistance given by Research IT and the use of the Computational Shared Facility at The University of Manchester, and the advice and support given by *EnginZyme*. CRF would also like to acknowledge Dr. Matthew Faulkner for his continued support and feedback during the bioinformatics pipeline and ColabAlign development processes.

