## Supporting Information document for "Identification of Diverse Cellulose Binding Domains using *in silico* Prioritisation and High-Throughput Screening"

[b]: EnginZyme, Tomtebodavägen 6, 171 65 Solna, Sweden

[c]: School of Chemical and Physical Sciences and Centre for Glycoscience, Keele University, Keele, Staffordshire, ST5 5BG, UK.

[d]: Manchester Institute of Biotechnology, Department of Chemistry, School of Natural Sciences, University of Manchester, 131 Princess Street, Manchester M1 7DN, UK.

Email:

;

;

;

;

### Supporting Information

**Table S1.** CBM genes selected for *in vitro* screening from structural dendrogram clustering.

| *Sequence Family* | *Cluster (CL)* | *GenBank Code* | *CBM-encoding region (AA)* | *Organism of Origin* | *Kingdom* | *Expression strain* | *Predicted substrate profile*[a] |
|---|---|---|---|---|---|---|---|
| 2 | 37 | QKX19027 | 2-91 | *Microbulbifer sp. YPW1* | *Bacteria* | *BL21(DE3)* | cellulose chitin xylan |
| 2 | 21 | AXI80431 | 263-345 | *Peterkaempfera bronchialis* | *Bacteria* | *BL21(DE3)* | cellulose chitin xylan |
| 2 | 12 | ABM79533 | 17-96 | *Prochlorococcus marinus str. MIT 9303* | *Bacteria* | *BL21(DE3)* | cellulose chitin xylan |
| 2 | 22 | ABV54446 | 371-462 | *Radopholus similis* | *Animalia* | *BL21(DE3)* | cellulose chitin xylan |
| 2 | 41 | QYY36597 | 692-786 | *Ruficoccus sp. ZRK36* | *Bacteria* | *BL21(DE3)* | cellulose chitin xylan |
| 3 | 13 | ADL50319 | 557-632 | *Clostridium cellulovorans 743B* | *Bacteria* | *BL21(DE3)* | cellulose chitin |
| 3 | 14 | ABX43721 | 775-857 | *Lachnoclostridium phytofermentans ISDg* | *Bacteria* | *BL21(DE3)* | cellulose chitin |
| 6 | 18 | AMJ62924 | 2094-2209 | *Bosea sp. PAMC 26642* | *Bacteria* | *BL21(DE3)* | β-glucan cellulose xylan |
| 6 | 36 | ARK10767 | 927-1065 | *Fibrella sp. ES10-3-2-2* | *Bacteria* | *BL21(DE3)* | β-glucan cellulose xylan |
| 6 | 33 | AJW82061 | 631-757 | *Flammeovirga sp. OC4* | *Bacteria* | *BL21(DE3)* | β-glucan cellulose xylan |
| 6 | 39 | UII22318 | 718-838 | *Fulvivirga ligni* | *Bacteria* | *BL21(DE3)* | β-glucan cellulose xylan |
| 6 | 34 | AHF89581 | 348-485 | *Opitutaceae bacterium TAV5* | *Bacteria* | *BL21(DE3)* | β-glucan cellulose xylan |
| 6 | 28 | AWV35627 | 1406-1515 | *Paenibacillus odorifer* | *Bacteria* | *BL21(DE3)* | β-glucan cellulose xylan |
| 6 | 40 | AIQ43223 | 651-779 | *Paenibacillus sp. FSL R5-0912* | *Bacteria* | *BL21(DE3)* | β-glucan cellulose xylan |
| 6 | 44 | BBD98785 | 345-464 | *Sphingobium amiense* | *Bacteria* | *BL21(DE3)* | β-glucan cellulose xylan |
| 6 | 19 | QIP11132 | 78-241 | *Spirosoma aureum* | *Bacteria* | *BL21(DE3)* | β-glucan cellulose xylan |
| 6 | 35 | AZP18224 | 848-965 | *Streptomyces aquilus* | *Bacteria* | *BL21(DE3)* | β-glucan cellulose xylan |
| 6 | 23 | AXL91867 | 1185-1299 | *Streptomyces sp. CB09001* | *Bacteria* | *BL21(DE3)* | β-glucan cellulose xylan |
| 6 | 38 | CAD5991209 | 698-828 | *Streptomyces sp. KY75* | *Bacteria* | *BL21(DE3)* | β-glucan cellulose xylan |
| 8 | 24 | BDT31982 | 259-398 | *Myxococcus sp. MH1* | *Bacteria* | *BL21(DE3)* | cellulose |
| 9 | 16 | UYO98914 | 886-1060 | *Oceanotoga sp. DSM 15011* | *Bacteria* | *BL21(DE3)* | cellulose |
| 9 | 15 | QUL53835 | 91-267 | *Paenibacillus tritici* | *Bacteria* | *BL21(DE3)* | cellulose |
| 9 | 11 | WKN31853 | 62-260 | *Poriferícola rhodea* | *Bacteria* | *BL21(DE3)* | cellulose |
| 16 | 29 | QDE02055 | 31-153 | *Erysipelothrix rhusiopathiae* | *Bacteria* | *BL21(DE3)* | cellulose glucomannan |
| 28 | 27 | AFI25187 | 570-760 | *Halalkalibacter akibai* | *Bacteria* | *BL21(DE3)* | cellulose |

| | | | | | | | |
|---|---|---|---|---|---|---|---|
| 30 | 25 | AFC28690 | 609-756 | *Paenibacillus mucilaginosus 3016* | *Bacteria* | *BL21(DE3)* | cellulose |
| 37 | 9 | ADU22937 | 381-444 | *Ruminococcus albus 7 = DSM 20455* | *Bacteria* | *BL21(DE3)* | cellulose<br>chitin<br>xylan |
| 44 | 26 | AUG56052 | 358-505 | *Acetivibrio saccincola* | *Bacteria* | *BL21(DE3)* | cellulose<br>xyloglucan |
| 46 | 8 | WFE61285 | 504-591 | *Micromonospora sp. WMMD712* | *Bacteria* | *BL21(DE3)* | cellulose |
| 46 | 7 | ASA25821 | 478-562 | *Paenibacillus donghaensis* | *Bacteria* | *BL21(DE3)* | cellulose |
| 49 | 6 | CAE6132321 | 530-608 | *Arabidopsis arenosa* | *Plantae* | *BL21(DE3)* | cellulose |
| 63 | 5 | BCB79642 | 88-161 | *Phytohabitans flavus* | *Bacteria* | *BL21(DE3)* | cellulose |
| 64 | 3 | WQG86787 | 825-897 | *Chitinophaga sancti* | *Bacteria* | *BL21(DE3)* | cellulose |
| 85 | 20 | QIG54529 | 75-196 | *Altererythrobacter sp. BO-6* | *Bacteria* | *BL21(DE3)* | β-glucan<br>β-mannan<br>cellulose<br>xylan |
| 1 | 2 | QKD61425 | 422-457 | *Fusarium oxysporum* | *Fungi* | *SHuffle T7* | *cellulose*<br>*chitin* |
| 2 | 31 | BCJ48281 | 354-448 | *Actinoplanes ianthinogenes* | *Bacteria* | *SHuffle T7* | *cellulose*<br>*chitin*<br>*xylan* |
| 2 | 42 | UQU62154 | 34-134 | *Couchioplanes caeruleus* | *Bacteria* | *SHuffle T7* | *cellulose*<br>*chitin*<br>*xylan* |
| 2 | 17 | USX50404 | 18-107 | *Lentzea sp. HUAS12* | *Bacteria* | *SHuffle T7* | *cellulose*<br>*chitin*<br>*xylan* |
| 2 | 43 | AXH93247 | 310-401 | *Micromonospora aurantiaca* | *Bacteria* | *SHuffle T7* | *cellulose*<br>*chitin*<br>*xylan* |
| 2 | 32 | SBT48127 | 377-477 | *Micromonospora auratinigra* | *Bacteria* | *SHuffle T7* | *cellulose*<br>*chitin*<br>*xylan* |
| 2 | 46 | CAN95714 | 46-140 | *Sorangium cellulosum So ce56* | *Bacteria* | *SHuffle T7* | *cellulose*<br>*chitin*<br>*xylan* |
| 2 | 47 | WAE72516 | 35-132 | *Streptomonospora nanhaiensis* | *Bacteria* | *SHuffle T7* | *cellulose*<br>*chitin*<br>*xylan* |
| 3 | 10 | AHL27899 | 492-569 | *Uncultured bacterium* | *Bacteria* | *SHuffle T7* | *cellulose*<br>*chitin* |
| 6 | 45 | WOO37091 | 321-442 | *Anaerocolumna sp. AGMB13020* | *Bacteria* | *SHuffle T7* | *β-glucan*<br>*cellulose*<br>*xylan* |
| 10 | 1 | ABD79918 | 409-439 | *Saccharophagus degradans* | *Bacteria* | *SHuffle T7* | *cellulose* |
| 16 | 30 | AJZ82552 | 47-161 | *Streptomyces sp. AgN23* | *Bacteria* | *SHuffle T7* | *cellulose*<br>*glucomannan* |
| 63 | 4 | UJV45240 | 134-207 | *Streptomyces sp. AMCC400023* | *Bacteria* | *SHuffle T7* | *cellulose* |

[a] Substrate binding profiles were predicted using dbCAN-sub (1)

**Table S2**. CBM families with known cellulose binding capabilities, not identified by dbCAN-sub.[a]

| | *Confirmed cellulosic substrates* | | |
|---|---|---|---|
| ***Sequence Family*** | ***Crystalline (≥ DP6)*** | ***Amorphous (DP4-6)*** | ***Derivatives*** |
| 4 | - | ASC (2)<br>DP6 (2) | HEC (3)<br>HPMC (4) |
| 5 | Avicel PH-101 (5,6) | Whatman No. 1 filter paper (6) | - |
| 11 | Avicel PH-101 (7) | ASC (7) | |
| 22 | Avicel PH-101 (8,9)<br>Avicel PH-105 (10)<br>Sigmacell 101 (9)<br>BMCC (9) | - | HEC (8,11)<br>MC (11) |
| 24 | MCC (25 μm average) (12) | - | - |
| 29 | - | Unspecified DP (13) | HEC (13) |
| 59 | Avicel (unspecified grade) (14) | - | - |
| 60 | - | - | CMC (15)<br>HEC (15)<br>HMC (15) |
| 65 | - | ASC (16) | HEC (16)<br>MC (17) |
| 72 | Avicel PH-101 (18) | ASC (18) | 2-HEC (18)<br>CMC (19)<br>MC (18) |
| 76 | - | - | HEC (20) |
| 78 | - | DP5+ (20) | HEC (20) |
| 79 | - | DP4+ (20) | HEC (20)<br>RC (20) |
| 80 | - | DP5+ (20) | HEC (20) |
| 81 | Avicel PH-101 (21) | Unspecified DP (21) | - |

[a] ASC is acid-swollen cellulose; BMC is ball-milled cellulose; BMCC is bacterial microcrystalline cellulose; CMC is carboxymethyl cellulose; DP is degree of polymerisation; HEC is hydroxyethyl cellulose; HPMC is hydroxypropylmethyl cellulose; MC is methyl cellulose; MCC is microcrystalline cellulose; RC is regenerated cellulose.

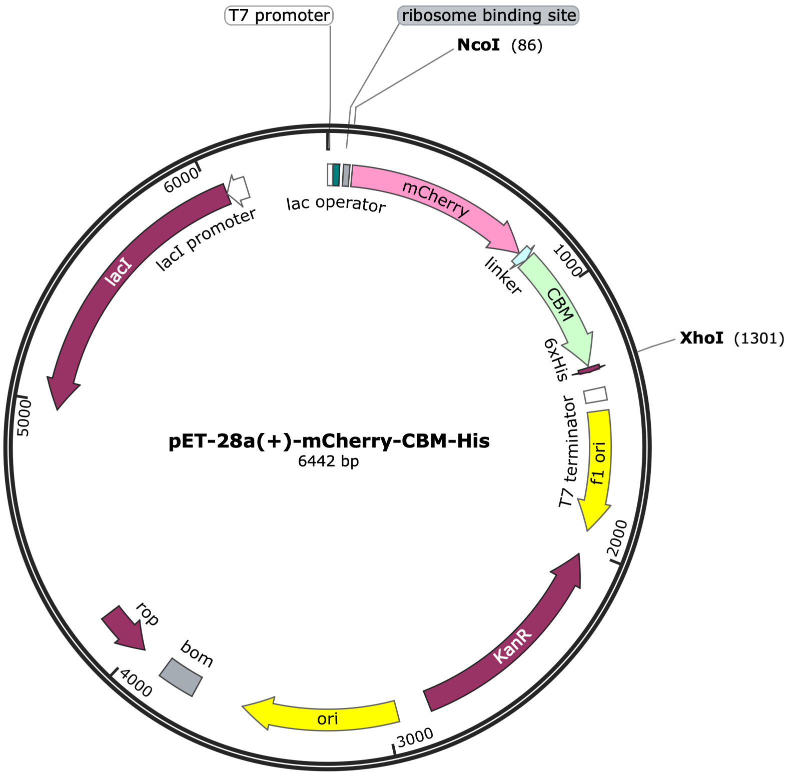


**Figure S1.** Example plasmid map for mCherry-CBM fusion expression in pET-28a(+).

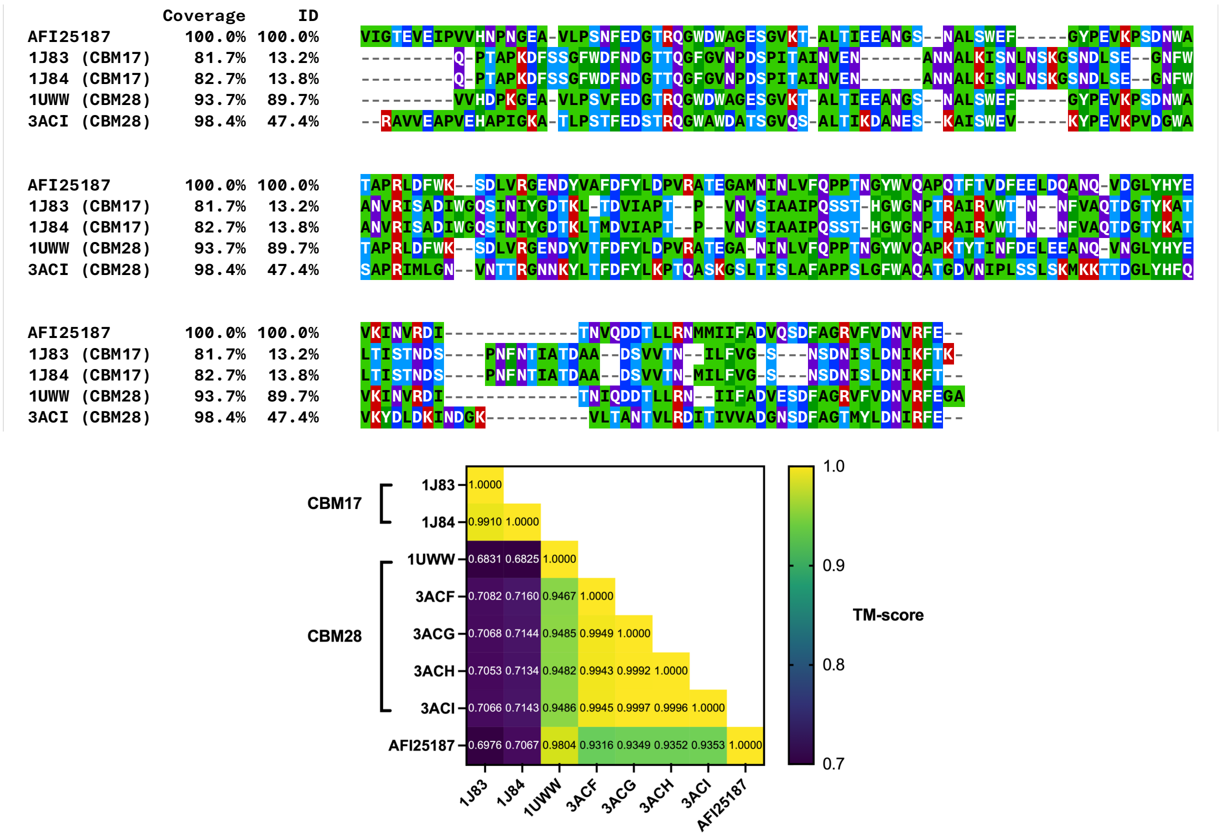


**Figure S2.** Structure-based sequence alignment (MUSTANG (22); MView (23)) and TM-score matrix (US-align (24)) comparing the AlphaFold structures of GenBank entry AFI25187 (average per-residue pLDDT: 95.52) and published structures for CBM17 (1J83, 1J84; (25)) and CBM28 (1UWW; (26); 3ACF, 3ACG, 3ACH, 3ACI; (27)). Sequences for 3ACF, 3ACG and 3ACH were excluded from the sequence alignment as they were identical to 3ACI. Sequence coverage and identity (ID) were calculated relative to AFI25187. Residue colouring follows the MView 'property' colour scheme: light green is hydrophobic; dark green is large hydrophobic; light blue is small alcohol; dark blue is negative; red is positive; purple is polar.

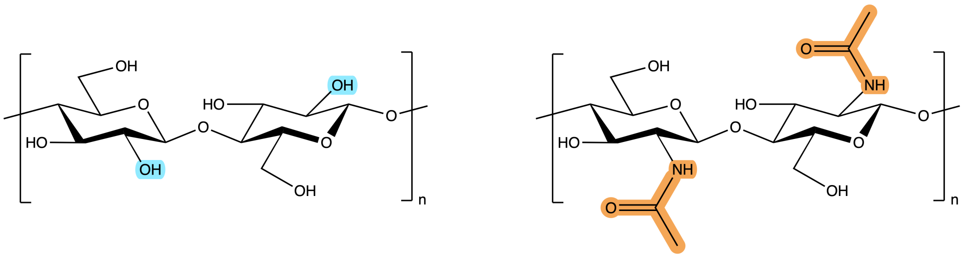


**Cellobiose** **Chitobiose**

**Figure S3.** Skeletal structures for the repeat units of cellulose (cellobiose) and chitin (chitobiose), highlighting the different moieties at the C2 position (hydroxyl for cellobiose; acetylamino for chitobiose).

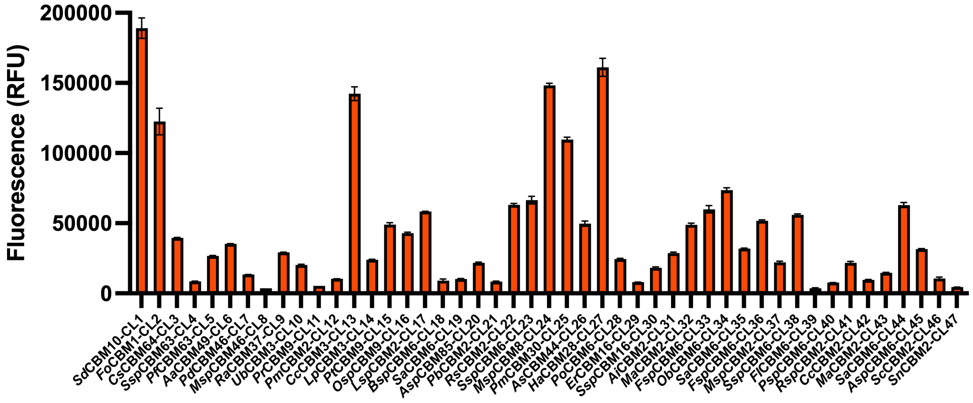


**Figure S4.** Raw soluble mCherry fluorescence data from preliminary expression trials.

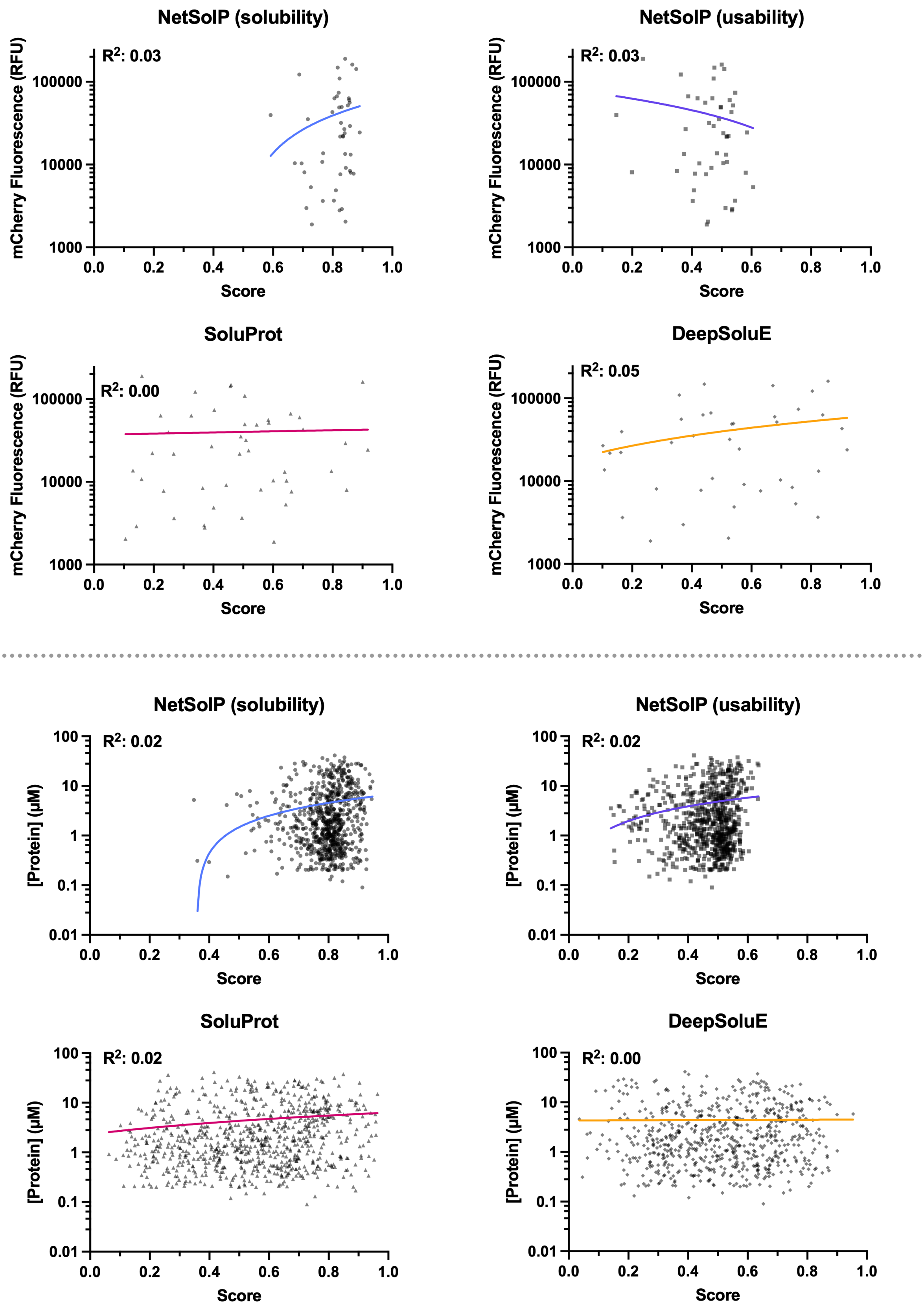


**Figure S5**. Correlation between *in silico* solubility and expression scores and observed expression levels for *Top:* preliminary expression data (**Figure S4**); and *Bottom:* expression data given by Rennison et al. (28).

### ColabAlign

**Pairwise Structural Alignment:** For a given set of protein structures, a list of all possible combinations of 2 structures (without replacement) is generated. This list is then evenly split into  sub-lists where  is the number of CPU cores assigned to the job. A subprocess is initialised for each of these sub-lists and pairwise structural alignments are performed using US-align in parallel. US-align outputs template modelling (TM) scores (29) for each alignment, normalised by the length of each of the protein structures. TM-scores are calculated using Eq. 1:

$$\text{TM-score} = \max\left[\frac{1}{L_{\text{target}}} \times \sum_{i}^{L_{\text{common}}} \frac{1}{1+\left(\frac{d_i}{\left(1.24\times\sqrt[3]{L_{\text{target}}-15}\right)-1.8}\right)^2}\right] \quad (1)$$

where  is the length of the target protein in residues;  is the number of residues that are present in the template and target proteins;  is the distance between the th pair of residues of the template and target proteins;  scales the magnitude of the final TM-score, making it length-independent.

**Dendrogram Generation:** The maximum of the TM-scores, corresponding with the score normalised by the smaller of the two structures, is stored in a lower triangular matrix. Once all pairwise alignments are complete, the values in the matrix are converted to distances for dendrogram generation using Eq. 2:

$$\text{distance} = 1 - \text{TM-score} \quad (2)$$

This inverts the values such that TM-scores trend towards 1.0 as distances trend towards 0.0. Distance values are used to calculate a dendrogram based on the global structural similarity between each structure using the UPGMA algorithm (30). This algorithm was selected as it is computationally simple relative to more modern algorithms, enabling quick processing of large datasets.

**Dendrogram Clustering:** Taxa on the dendrogram are clustered with TreeCluster (31) with a default distance threshold of 0.25 (equivalent to a TM-score of 0.75). For each cluster in the TreeCluster output file, a corresponding directory is created, into which the associated structure files are copied. A temporary reference model is selected from the cluster by generating a submatrix containing the TM-scores of only the members of the cluster, then calculating the average TM-score for each model and selecting the model with the highest average. The coordinates of the non-reference models are then superimposed over the reference model using the transformation matrix output from US-align for that pair using Eq. 3:

$$T_{\text{forward}} = [t_{\text{forward}}\, r_{\text{forward}}] = \begin{bmatrix} t_0 & u_{00} & u_{01} & u_{02} \\ t_1 & u_{10} & u_{11} & u_{12} \\ t_2 & u_{20} & u_{21} & u_{22} \end{bmatrix}, \text{ where } T_{\text{forward}} \in \mathbb{R}^{3\times4}, \quad (3)$$

where  represents the transformation matrix;  is the translation vector;  is the rotation matrix. The transformation matrix is applied to the 3D coordinates of each atom of the non-reference protein using Eq. 4:

$$\begin{bmatrix} x_2 \\ y_2 \\ z_2 \end{bmatrix} = \left( r_{\text{forward}} \cdot \begin{bmatrix} x_1 \\ y_1 \\ z_1 \end{bmatrix} \right) + t_{\text{forward}}, \text{ where } \begin{bmatrix} x_1 \\ y_1 \\ z_1 \end{bmatrix}, \begin{bmatrix} x_2 \\ y_2 \\ z_2 \end{bmatrix} \in \mathbb{R}^3, \quad (4)$$

where , , and are Cartesian coordinates; 1 is the start coordinates; 2 is the end coordinates. In cases where the reference model was transformed onto another model as part of the original TM-score matrix calculations, the transformation matrix is reversed using Eq. 5:

$$T_{\text{reverse}} = -\left(r_{\text{forward}}^{\mathrm{T}} \cdot t_{\text{forward}}\right),\ \text{where}\ r_{\text{forward}}^{\mathrm{T}} = \begin{bmatrix} u_{00} & u_{10} & u_{20} \\ u_{01} & u_{11} & u_{21} \\ u_{02} & u_{12} & u_{22} \end{bmatrix}, \tag{5}$$

where  is the transposition of . Since cluster '-1' contains models which could not be clustered, no alignment operations are performed.

**Cluster representative selection:** Structure-informed sequence alignments for each cluster, excluding cluster '-1', are calculated using MUSTANG (22). Each MUSTANG job runs in a separate subprocess. The resulting aligned FASTA for each cluster is analysed using MView (23) to generate consensus sequences at 60, 70, 80, 90 and 100% identity. The aligned amino acid sequence for each model in the cluster is then compared against the 60% identity consensus sequence for the cluster. MView uses an expanded character set to denote where non-identical amino acids with similar properties have been aligned (**Table S3**). Each alignment is scored on a per-residue basis, where the maximum score for the alignment is its length, including gap characters. This is identical for each sequence in a given cluster. The cumulative score for each alignment is incremented by 1 in cases where a gap character is present in both sequences; identical residues are present; or where the current residue of the query sequence falls under the same group as the MView character in the consensus sequence. The sequence with the highest score of the cluster is selected as the representative for that cluster.

**Table S3.** MView expanded character set for sequence alignments.

| *Group* | *MView character* | *1 letter amino acid codes* |
|---|---|---|
| *Alcohol* | o | S, T |
| *Aliphatic* | l | I, L, V |
| *Aromatic* | a | F, H, W, Y |
| *Charged* | c | D, E, H, K, R |
| *Hydrophobic* | h | A, C, F, G, H, I, K, L, M, R, T, V, W, Y |
| *Negative* | - | D, E |
| *Polar* | p | C, D, E, H, K, N, Q, R, S, T |
| *Positive* | + | H, K, R |
| *Small* | s | A, C, D, G, N, P, S, T, V |
| *Tiny* | u | A, G, S |
| *Turn-like* | t | A, C, D, E, G, H, K, N, Q, R, S, T |
| *Stop* | * | * |
